# The AMPK activator A-769662 inhibits human TASK3 potassium channels in an AMPK-independent manner

**DOI:** 10.1101/2022.05.24.493214

**Authors:** Esraa A. Said, Ryan W. Lewis, Mark L. Dallas, Chris Peers, Fiona A. Ross, D. Grahame Hardie, A. Mark Evans

**Author notes:** **CORRESPONDING AUTHOR:** A. Mark Evans, Centre for Discovery Brain Sciences, College of Medicine and Veterinary Medicine, Hugh Robson Building, University of Edinburgh, Edinburgh, EH8 9XD, UK. Esraa Said is also an Assistant Lecturer at the Department of Pharmacology and Toxicology, Faculty of Pharmacy, Alexandria University. **AUTHOR CONTRIBUTIONS** A.M.E. conceived of this study. A.M.E. and E.S. designed the experiments presented here. A.M.E. and E.S. wrote this manuscript and made all Figures. E.S., R.W.L. and M.L.D. carried out patch-clamp electrophysiology and completed data acquisition. A.M.E. and E.S. analysed the data. D.G.H. and F.A.F. carried out TASK-3 phosphorylation and provided HEK293 cells expressing the AMPK β-1 S108A mutant. C.P. oversaw the experiments completed by M.L.D. and sourced the HEK293 cell line that stably expressed human TASK-3.

## Abstract

Heteromeric TASK1/3 channels play a fundamental role in oxygen-sensing by carotid body type 1 cells, where hypoxia-induced inhibition of TASK3 and/or TASK1/3 potassium currents leads to depolarisation, voltage-gated calcium entry, exocytotic transmitter release and increases in carotid body afferent input responses that initiate corrective changes in breathing patterns. However, the mechanism by which hypoxia leads to TASK-1/3 channel inhibition is still debated. It had been proposed that the AMP-activated protein kinase (AMPK) might directly phosphorylate and inhibit TASK channels, in particular TASK-3, although subsequent studies on rat type I cells argued against this view. Here we report on the effects of novel, highly selective AMPK activators on recombinant human TASK-3 potassium channels. Sequence alignment identified an AMPK recognition motif in TASK-3, but not TASK-1, with Ser^55^ representing a potential site for AMPK-dependent phosphorylation in TASK-3. However, neither of the AMPK activators, AICAR or MK-8722, caused a significant reduction of human TASK-3 current amplitude. By contrast, high concentrations of the AMPK activator A-769662 (100-500 µM) inhibited human TASK-3 currents in a concentration-dependent manner. Importantly, A-769662 (300 µM) also inhibited human TASK-3 channels in HEK293 cells that stably over-expressed an AMPK-β1 subunit mutant (S108A) that renders AMPK insensitive to activators binding the Allosteric Drug and Metabolite (ADaM) site, such as A-769662. We therefore identify A-769662 as a novel human TASK-3 channel inhibitor and provide conclusive evidence that AMPK does not regulate TASK-3 channel currents.

## INTRODUCTION

Two pore domain K^+^ channels (K2P) are a large family of K^+^ channels that are widely distributed in numerous tissue types (1). K2P channels are voltage- and time-independent channels that are constitutively open over the entire range of physiological potentials and hence are also known as leak or background potassium channels, exerting major influences on cell membrane potential and excitability (2-4). Nevertheless, K2P channels have been shown to provide much more than just a leak current (5). They are tightly regulated by various stimuli, such as pH, temperature, membrane stretch, G proteins, fatty acids, and protein kinases. Indeed, their important physiological roles and multiple means of regulation have led to intensive investigations of their therapeutic potential in cardiovascular and neurological disorders, as well as in cancer (6, 7).

Within the K2P family, a sub-group exists that was originally recognised as being particularly sensitive to extracellular pH: the TASK channels (where TASK represents <u>T</u>andem (or <u>T</u>wo) pore domain <u>A</u>cid <u>S</u>ensitive <u>K</u>^+^ channel). This subfamily comprises TASK-1 (other names, KCNK3 and K2P3.1), TASK-3 (KCNK9 and K2P9.1), and TASK-5 (KCNK15 and K2P15.1). These TASK isoforms show different pH-sensitivities. TASK-1 exhibits higher pH sensitivity than TASK-3 (circa 0.5 pH units), whereas that of TASK-1/TASK-3 heteromeric channels is intermediate (8). Nevertheless, identifying TASK channels only based on their extracellular pH sensitivity does not distinguish them from other K2P channels, such as TWIK-1, TALK-1, and TALK-2, which show similar pH sensitivity and biophysical characteristics. Therefore, other TASK-selective pharmacological criteria have to be used, such as oxygen sensitivity (in specific cell types), sensitivity to volatile anaesthetics and the use of TASK isoform-selective blockers, including anandamide and ruthenium red. Native TASK-like currents have been identified in many cell types, including carotid body type 1 cells and hippocampal pyramidal neurons (9). Shortly before the molecular identification of TASK channels, Buckler (10) characterised a “leak” K^+^ conductance in type I cells of the rat carotid body, which was not only TASK-like in its sensitivity to extracellular pH (being inhibited by acidification), but was also inhibited by hypoxia. This K^+^ current, and the channels which gave rise to it, was of particular importance in the field as it provided a direct means by which these cells could be responsive to either hypoxia or acidosis, the two primary physiological stimuli of the carotid body. Although both TASK-1 and TASK-3 isoforms are present in carotid body type 1 cells, the dominant oxygen-sensitive leak current has been shown to be carried by TASK-1/3 heteromeric channels (11, 12). Whilst other K^+^ channels have also been implicated in hypoxia and acid sensing in type 1 cells (13-16), TASK channels have been shown to play a central role in chemotransduction. Briefly, TASK channel inhibition leads to depolarization, voltage-gated Ca2+ entry, exocytotic transmitter release and increased carotid body afferent input responses that initiate corrective changes in breathing patterns (17).

A fundamental question concerning oxygen-sensing by K^+^ channels in general, and by TASK channels is the mechanism by which hypoxia leads to their inhibition. We previously proposed that the AMP-activated protein kinase (AMPK) may mediate the oxygen sensitivity of carotid body type 1 cells via direct phosphorylation and inhibition of TASK channels based primarily on the action of the AMPK activator AICAR (18-20). However, there is controversy about this hypothesis because others have shown that two different AMPK activators, AICAR and A769662, are without effect on TASK-1/3 currents in rat carotid body type I cells (21, 22), countering previous contrary reports on the effects of AICAR on rat type I cell TASK currents and carotid body afferent discharge (19). Contrary to this view, a preliminary study carried out on recombinant human (h) TASK-3 channels showed hTASK-3 channel inhibition by A769662 (20). Therefore, regulation of TASK-3 channels by AMPK remains open to debate.

Here we have revisited the potential modulation of hTASK-3 channels by AMPK, using the AMPK activator MK-8722 which, like A-769662, binds at the Allosteric Drug and Metabolite (ADaM) site (23, 24). MK-8722 is a more potent activator than A-769662, and (unlike the latter which is β1-specific) activates AMPK complexes containing the β1 or β2 subunit isoforms. Our results confirm beyond doubt that AMPK does not contribute to the functional modulation of recombinant human TASK-3 channels despite the presence of a potential AMPK recognition motif. Moreover, we identify A769662 as a novel concentration-dependent hTASK3 channel inhibitor that acts independent of AMPK.

## METHODS

### Generation of TASK-3 expressing cell lines

HEK293 cells were cultured in modified Eagle’s medium (MEM) with Earle’s salts and L-glutamine, supplemented with 9% (v/v) fetal calf serum (Globepharm, Esher, Surrey, UK), 1% (v/v) non-essential amino acids, 50 μg/ml gentamicin, 100 units/ml penicillin G, 100 μg/ml streptomycin and 0.25 μg/ml amphotericin in a humidified atmosphere of air/CO_2_ (19:1) at 37°C. All cell culture reagents were purchased from Gibco-BRL (Paisley, UK) unless otherwise stated.

HEK-293 cells stably expressing FLAG-tagged AMPK-β1 with either the wild type sequence or an S108A mutation have been described previously (25); in these cells the FLAG-tagged β1 subunit effectively replaces the endogenous β1 and β2 subunits. Full length cDNAs encoding human TASK-3 (hTASK-3) channels, a kind gift from Dr. S. A. N. Goldstein (Department of Pediatrics and Institute for Molecular Pediatric Sciences, Pritzker School of Medicine, University of Chicago), were originally subcloned into the mammalian expression vector pMAX(+) via Xba I/Xma I restriction site combinations.

To generate HEK293 cell lines stably expressing wild-type hTASK-3, cells were transfected with pMAX/hTASK-3 constructs, using the PolyFect transfection reagent (Qiagen, Hybaid Ltd, Teddington, UK) according to manufacturer’s instructions. Stable HEK293 cell lines were achieved by antibiotic selection with G-418 (1mg/ml, Gibco-BRL, Paisley, UK) added to the medium 3 days after transfection. Selection was applied for 4 weeks (media changed every 4-5 days), after which time individual colonies were picked and seeded in T25 flasks and allowed to reach confluence. They were then transferred to T75 flasks for further culture and electrophysiological screening. Cells were harvested from culture flasks by trypsinization and plated onto coverslips 24-48h before use in electrophysiological studies.

Transfection of hTASK-3 channels was considered successful if the currents elicited by the whole-cell voltage ramp protocol (see below) were: a) significantly (> 4-fold) larger than untransfected HEK293 cell K^+^ currents; b) reliably described by the Goldman Hodgkin and Katz (GHK) equation and; c) activated by pH 8.4 and inhibited by pH 6.4 (and the pH-sensitive currents demonstrated GHK rectification). In addition, hTASK-3 was inhibited by ruthenium red, as previously described [18].

### Cell transfection

human TASK-3 was overexpressed by transient transfection (Lipofectamine 2000 (Invitrogen)) into a HEK293 cell line stably expressing AMPK-β-1S108A mutant subunit, that were seeded on 100-mm dishes. At 48 h post-transfection, cells were washed with bath solution and used for electrophysiology.

### Electrophysiology

Whole-cell patch-clamp recordings were acquired from HEK293 cells stably expressing either the hTASK-3, or HEK 293 cells stably expressing the AMPK-β1 S108 A mutant that were transiently expressed wild-type hTASK-3. Coverslips with attached cells were transferred to a continuously perfused recording chamber (perfusion rate 3-5 ml/min, volume ca 200μl) mounted on the stage of an inverted microscope (Zeiss, Axioscope). Cells were perfused with a solution containing (in mM): 135 NaCl, 5 KCl, 1.2 MgCl_2_, 5 HEPES, 2.5 CaCl_2_, 10 D-glucose (pH 7.4 with KOH). Patch electrodes (resistance 4-7MΩ) were filled with intracellular solution consisting of (in mM): 10 NaCl, 117 KCl, 2 MgCl_2_, 11 HEPES, 11 EGTA, 1 CaCl_2_, 2 Na_2_ATP (pH 7.2 with KOH).

Outward K^+^ currents were recorded at 37°C (unless otherwise stated) using standard ramp protocols; cells were voltage-clamped at a holding potential of –70mV, then stepped to -100mV and a voltage ramp (500ms duration) immediately applied from -100 mV to +60mV. Data were acquired and digitized via a Digidata 1322A in combination with an Axopatch 200B amplifier and Clampex 10 software (Molecular Devices, Foster City, CA). Currents were sampled at 2kHz and low-pass filtered at 1kHz. Offline data analysis was conducted using Clampfit 10 software (Molecular Devices, Foster City, CA.).

### Statistical analysis

Statistical analysis was completed using the paired Student’s *t-test*, One-way ANOVA followed by Bonferroni’s post-hoc test, or Two-way ANOVA followed by Tukey’s post-hoc test. Differences were considered significant when P<0.05. All values stated are as mean ± SEM.

### Drugs and chemicals

All chemicals were obtained from Sigma-Aldrich (Poole, UK) except MK-8722, which was synthesized by Dr Natalia Shpiro (University of Dundee) using published methods (26). MK-8722 and A769662 were applied to cells by direct bath application.

## RESULTS

### Recombinant hTASK-3 channels have an AMPK recognition motif

To examine whether hTASK-1 or hTASK-3 might be regulated by AMPK through direct phosphorylation, we examined their amino acid sequence for a motif which might be comparable to those found in recognised AMPK substrates. We identified no recognition motifs in hTASK-1, although we did identify a possible AMPK recognition motif in hTASK-3. Figure 1*A* illustrates an optimal motif, with idealised hydrophobic (H) and basic (B) residues surrounding a candidate serine residue (phospho site). Sequence alignment suggested that Ser^55^ in hTASK-3 represented a possible candidate site for AMPK phosphorylation, although it is not ideal because it lacks a bulky hydrophobic residue at the P-5 position, while the tyrosine at the P+4 position may also be sub-optimal (27). Consistent with this, we were unable to demonstrate 32^P^ incorporation from radiolabelled ATP using purified AMPK as we have done successfully for other ion channels regulated by AMPK, such as K_V_1.5, K_V_2.1 and K_Ca_1.1 (28-31).

**Figure 1.**
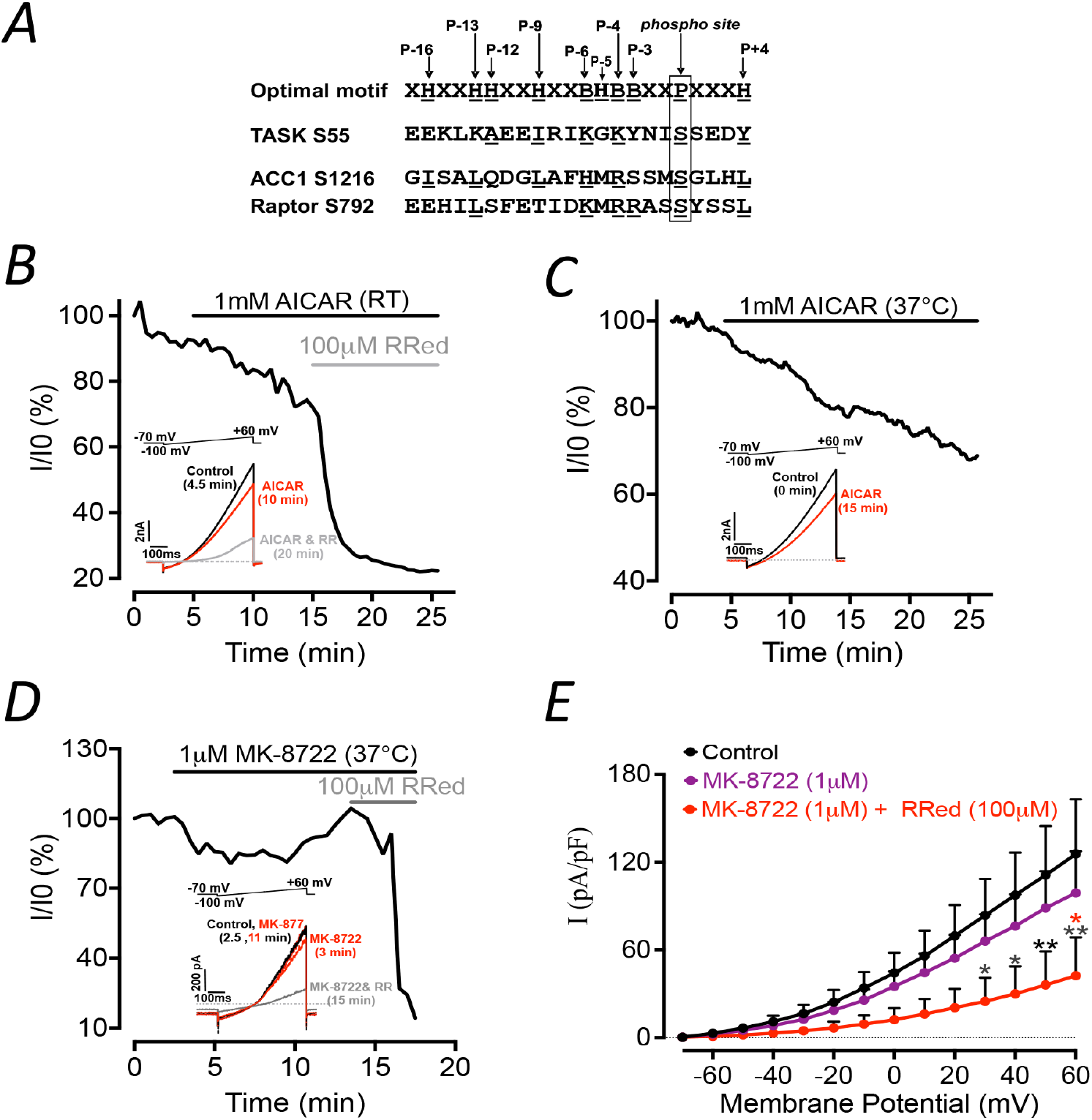
Different AMPK activators have no effect on hTASK-3 activity. *A, A*, Alignment of putative motifs for AMPK phosphorylation. Upper sequence is hypothetical idealised motif, and below are shown sequences for hTASK-3, acetyl-CoA carboxylase (ACC1) and Raptor. Phosphorylation sites are boxed; basic/hydrophobic residues likely to be involved in recognition are marked by underlining. *B*, Time course of hTASK-3 currents evoked by successive ramp depolarizations following extracellular application of 1mM AICAR (black line) and 100 μM Ruthenium Red (RR) (grey line) at room temperature (RT); or *C*, 1mM AICAR at 37°C (black line); or *D*, 1 μM MK-8722 (black line) and 100 μM RR (grey line) at 37°C; measurements taken at +60mV. Insets, hTASK-3 currents as indicated (*B)*, before (t = 0) and at 10 or 20 min after extracellular application of 1mM AICAR alone or combined with 100 μM RR at RT, respectively; (*C)*, before (0 min) and at 15 min after extracellular application of 1mM AICAR at 37°C; (*D)*, before (2.5 min) and at 3 and 11, or 15 min after extracellular application of 1 μM MK-8722 alone or combined with 100 μM RR at 37°C, respectively. *C*, I–V relationships for hTASK-3 current recorded before (control), and after extracellular application of 1 μM MK-8722, and after extracellular application of 100 μM RR in the continued presence of MK-8722; measurements taken at test potential of +60mV. Results are expressed as mean ± SEM, n = 4–5. Statistical analysis was carried by two-way ANOVA followed by Tukey’s post-hoc test, *denotes significance from control group and * denotes significance from the MK-8722 group, *P < 0.05, **P < 0.01.

### AICAR and MK-8722 have no effect on recombinant hTASK-3 channels

We next assessed the effect of AMPK activation on hTASK-3 currents using either AICAR or MK-8722. Extracellular application of 1 mM AICAR for 10-20 minutes at room temperature caused a slight reduction in hTASK-3 currents (Fig. 1*B*) but this decline was not significant compared to control cells after 20 min of AICAR application (Fig.4*A*). To confirm the presence of hTASK-3 channels, 100µM ruthenium red (RR) was bath-applied following AICAR and caused a prominent and rapid reduction of the currents (Fig. 1*B*). As AICAR is a prodrug that is activately converted to ZMP by cellular uptake and metabolism (32), we repeated the same experiment at 37°C. However, once again no significant hTASK-3 channel inhibition was observed beyond rundown seen in control cells, as exemplified in Fig. 1*C*. To investigate this further, we examined the effects of MK-8722, one of the most potent and selective pan-β AMPK activators currently available (23, 24). As with AICAR, direct bath application of 1µM MK-8722 failed to inhibit hTASK-3 currents (Fig. 1*D* and 1*E*, Fig. 4*A*), whereas subsequent addition of RR virtually abolished hTASK-3 currents (Fig. 1*D* and 1*E*).

### A-79662 inhibits hTASK-3 currents in a concentration-dependent and AMPK-independent manner

The study of Kim and colleagues (22) showed that A-769662 (100µM) had no effect on rat (r) TASK-3 channel activity in rat carotid body glomus cells. We therefore carried out similar studies using hTASK-3 channels stably expressed in HEK293 cells. Much like AICAR and MK-8722, we found that 100 µM A-769662 failed to cause a significant reduction of hTASK-3 (Fig. 2*Aa*, 2*Bb* and 4*A*) despite a previous report to the contrary (20). Surprisingly, however, higher concentrations of A-769662 (300 µM and 500 µM) caused rapid and marked inhibition of hTASK-3 currents, as shown in Fig. 2*Ab* and *c*, and 2*Bb* and *c* respectively; and Fig. 4*A*. As shown in Fig. 2*C*, 500 µM A-769662 caused a significant reduction of hTASK-3 current density as compared to paired control cells. It was also consistently observed that currents were transiently enhanced immediately following exchange of the normal/control bath solution for one containing A-769662, before being inhibited by this compound. Furthermore, A-769662-induced inhibition of hTASK-3 was concentration-dependent as exemplified in Fig 2*Ad* and 2*Bd*. Given that neither AICAR nor MK-8722 showed a significant effect on hTASK-3 channels, we attribute the observed A-769662-induced inhibition of hTASK-3 to be an off-target, AMPK-independent effect. To assess this possibility, we tested the effect of A-769662 (300 µM) on hTASK-3 channels expressed in HEK 293 cells that stably over express a FLAG-tagged AMPK-β1 wild type or S108A mutant; by competing for binding to the β and β subunits, the FLAG-tagged β1 subunits essentially replace the endogenous β1 and β2 subunits (25). A-796662 binding to the ADaM site on AMPK, in the cleft between the N-lobe of the kinase domain on the β subunit and the Carbohydrate-Binding Module on the β subunit (β-CBM) (33, 34). Phosphorylation of Ser108 in the β-CBM by autophosphorylation or by upstream kinases stabilises the ADaM site by interacting with two lysine residues in the kinase domain N-lobe, and an S108A mutation essentially abolishes the effects of ADaM site activators (25, 33-35). Therefore, if the inhibition of hTASK-3 channels by A-769662 is AMPK-independent, we would expect to observe similar levels of inhibition despite expression of the S108A mutant. Consistent with this, A-769662 (300 µM) significantly reduced hTASK-3 currents even in cells expressing an AMPK-β1 S108A mutant (Fig. 3 and Fig. 4*B*).

**Figure 2.**
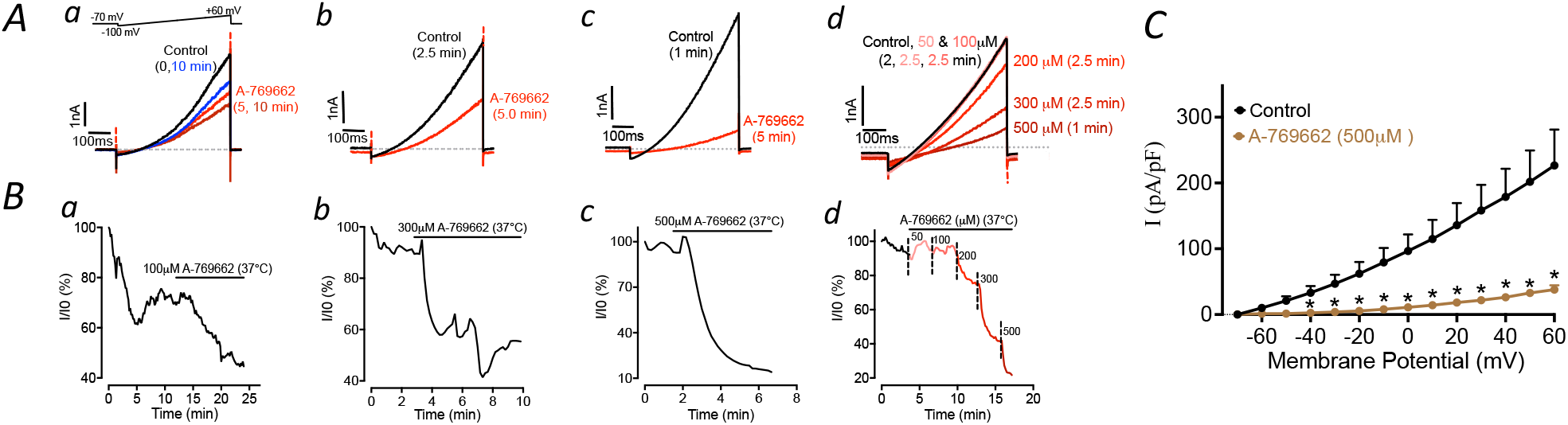
A-769662 inhibits hTASK-3 in a concentration-dependent manner. *A*, Example records of hTASK-3 currents at 37°C before (control) and after extracellular application of A-769662 at 100 μM (*a*), 300 μM (*b*), 500 μM (*c*), and in gradually increasing cumulative concentrations (from 50 to 500 μM, *d*) are shown. *B*, Time course for reduction of hTASK-3 currents following extracellular application of A-769662 at 100 μM (*a*), 300 μM (*b*), 500 μM (*c*), and in gradually increasing cumulative concentrations (from 50 to 500 μM, *d*) at 37°C. *C, I*–V relationships for hTASK-3 activity recorded before (control), and after extracellular application of 500 μM A-769662; measurements taken at +60mV. Results are expressed as mean ± SEM, n = 5. Statistical analysis was carried by paired t-test. *P < 0.05, **P < 0.01.

**Figure 3.**
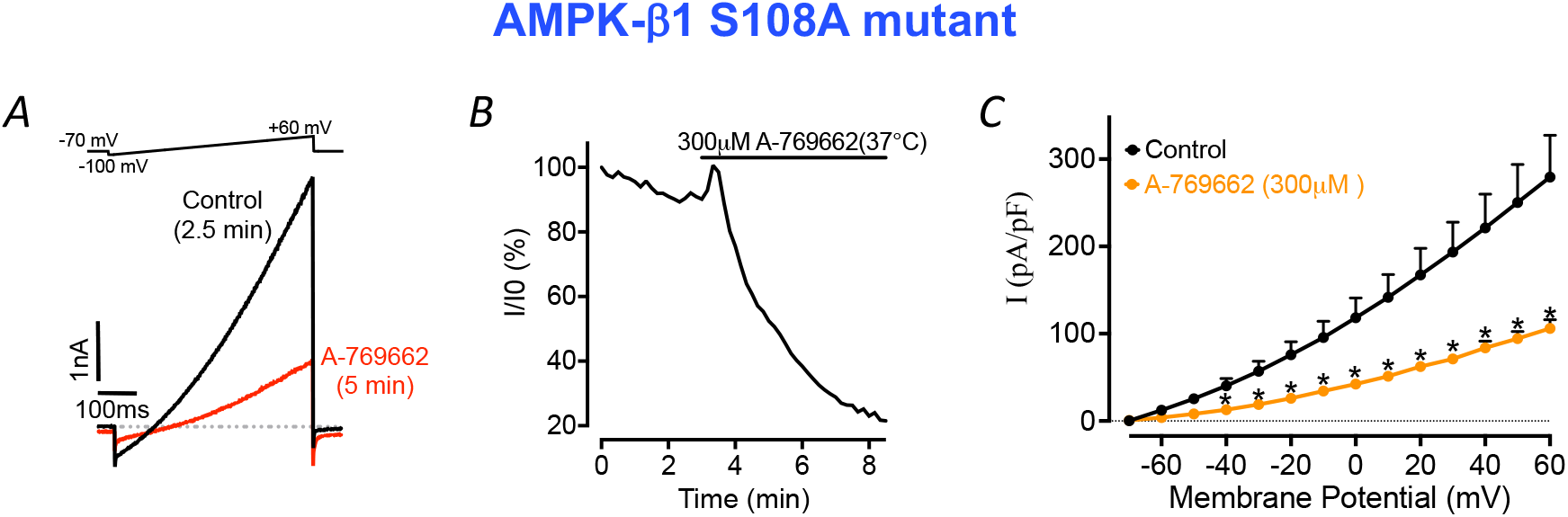
A-769662 inhibits hTASK-3 in an AMPK-independent manner. *A*, Example records of hTASK-3 currents transiently over-expressed in HEK293 cells, stably expressing AMPK β-1 S108A mutant, before (control) and after extracellular application of 300 μM A-769662 at 37°C. *B*, Time course for reduction of hTASK-3 currents following extracellular application of 300 μM A-769662. *C, I*–V relationships for hTASK-3 recorded before (control), and after extracellular application of 300 μM A-769662; measurements taken at test potential of +60mV. Results are expressed as mean ± SEM, n = 6. Statistical analysis was carried by paired t-test. *P < 0.05.

**Figure 4.**
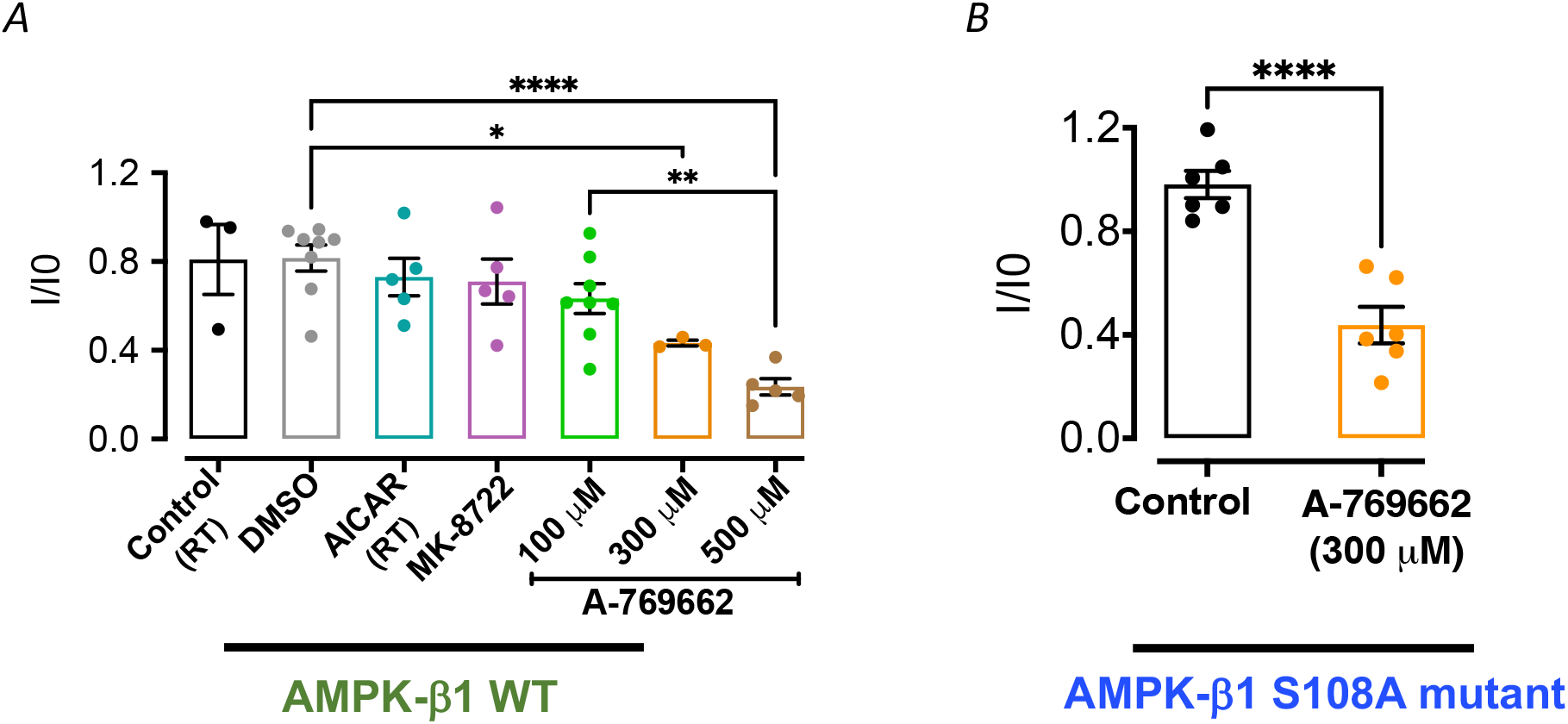
Comparison of the change in hTASK-3 current induced by different AMPK activators in HEK293 cells. Bar graphs illustrating normalized hTASK-3 currents stably expressed in AMPK-β1 wild-type (WT)-stably expressed HEK293 cells (*A*); and transiently co-expressed in AMPK β-1 S108A mutant-stably expressed HEK293 cells (*B*) in the presence of various AMPK activators at 37°C (unless otherwise stated); measurements taken at +60mV. Results are expressed as mean ± SEM, n =3-8. Statistical analysis was carried by one-way ANOVA followed by Bonferroni’s post-hoc test (*A*); and unpaired t-test (*B*). DMSO is the vehicle for A-769962 and MK-8722 and provides the relevant control. \**P* < 0.05, \*\**P* < 0.01, \*\*\**P* < 0.001 and \*\*\*\**P* < 0.0001.

## DISCUSSION

Even though we identified a possible AMPK recognition motif within human (h) TASK-3, but not hTASK-1, the present study suggests that AMPK does not regulate recombinant hTASK-3 potassium channels. Interestingly, however, we identified that A769662, an AMPK agonist, inhibits hTASK-3 in an AMPK-independent manner.

We found no significant phosphorylation of hTASK-3 by AMPK in cell-free assays. Furthermore, we found no evidence of hTASK-3 inhibition by two compounds that activate AMPK by different mechanisms, i.e., AICAR, a pro-drug that is metabolised to the AMP mimetic ZMP (32), and MK8722, a highly potent and selective activator of all AMPK isoforms that binds at the ADaM site (23, 24). These findings are in accordance with previous studies on the effects of two AMPK activators, namely AICAR and A769662, on TASK currents in rat carotid body type I cells (21, 22). When taken together, therefore, our study and those of others provides strong support for the view that AMPK does not phosphorylate or functionally modulate the hypoxia-responsive TASK-3 K^+^ channels, and refutes previous contrary reports on the effects of AICAR on rat type I cell leak currents (19) and AMPK-dependent inhibition of hTASK-3 (20).

Surprisingly, we found that extracellular application of A-769662 rapidly inhibits human TASK-3 channels in a concentration-dependent manner (≥300µM). As mentioned previously, A-796662 is an allosteric activator of AMPK through binding to the ADaM site. Therefore, we transiently expressed hTASK-3 in HEK293 cells that stably overexpressed a non-phosphorylatable AMPK β-1 S108A mutant that has been shown to block AMPK regulation compounds that bind to the ADaM site (25, 33-35). In HEK239 cells that stably over-expressed AMPK-β-1 S108A we observed a similar level of inhibition by high concentrations of A-769662 to that seen for cells expressing the wild-type. In short, A769662 inhibits hTASK-3 channels in an off-target, AMPK-independent manner, adding to previous reports of off-target inhibition of the sodium-potassium ATPase (36) and glucose uptake (37). Contrary to this, A-769662 has been shown to be without effect on TASK-1/3 currents in rat carotid body type I cells (21, 22) and therefore may exhibit selectivity for hTASK-3 over rat TASK-3 species dependency (human vs rat). This may have implications for future drug development against certain cardiovascular and neurological disorders where TASK channels have been implicated (6, 7).

## Author Agreement

This is to certify that all authors have seen and approved the final version of the manuscript being submitted

## References

1. Bayliss DA, Barrett PQ. Emerging roles for two-pore-domain potassium channels and their potential therapeutic impact. Trends Pharmacol Sci. 2008;29(11):566–75.

2. Natale AM, Deal PE, Minor DL. Structural Insights into the Mechanisms and Pharmacology of K2P Potassium Channels. Journal of Molecular Biology. 2021;433(17):166995.

3. Ilan N, Goldstein SA. Kcnkø: single, cloned potassium leak channels are multi-ion pores. Biophys J. 2001;80(1):241–53.

4. Mathie A, Al-Moubarak E, Veale EL. Gating of two pore domain potassium channels. J Physiol. 2010;588(Pt 17):3149–56.

5. Renigunta V, Schlichthörl G, Daut J. Much more than a leak: structure and function of K_2_p-channels. Pflugers Arch. 2015;467(5):867–94.

6. Bittner S, Budde T, Wiendl H, Meuth SG. From the background to the spotlight: TASK channels in pathological conditions. Brain Pathol. 2010;20(6):999–1009.

7. Es-Salah-Lamoureux Z, Steele DF, Fedida D. Research into the therapeutic roles of two-pore-domain potassium channels. Trends in Pharmacological Sciences. 2010;31(12):587–95.

8. Czirjak G, Enyedi P. Formation of functional heterodimers between the TASK-1 and TASK-3 two-pore domain potassium channel subunits. J Biol Chem. 2002;277(7):5426–32.

9. Lotshaw DP. Biophysical, pharmacological, and functional characteristics of cloned and native mammalian two-pore domain K+ channels. Cell Biochem Biophys. 2007;47(2):209–56.

10. Buckler KJ. A novel oxygen-sensitive potassium current in rat carotid body type I cells. J Physiol. 1997;498 (Pt 3)(Pt 3):649–62.

11. Kim D, Cavanaugh EJ, Kim I, Carroll JL. Heteromeric TASK-1/TASK-3 is the major oxygen-sensitive background K+ channel in rat carotid body glomus cells. J Physiol. 2009;587(Pt 12):2963–75.

12. Buckler KJ. TASK channels in arterial chemoreceptors and their role in oxygen and acid sensing. Pflugers Archiv : European journal of physiology. 2015;467(5):1013–25.

13. Hescheler J, Delpiano MA, Acker H, Pietruschka F. Ionic currents on type-I cells of the rabbit carotid body measured by voltage-clamp experiments and the effect of hypoxia. Brain Res. 1989;486(1):79–88.

14. Peers C. Hypoxic suppression of K+ currents in type I carotid body cells: selective effect on the Ca2(+)-activated K+ current. Neurosci Lett. 1990;119(2):253–6.

15. Peers C. Effect of lowered extracellular pH on Ca2(+)-dependent K+ currents in type I cells from the neonatal rat carotid body. J Physiol. 1990;422:381–95.

16. Patel AJ, Honoré E. Molecular physiology of oxygen-sensitive potassium channels. European Respiratory Journal. 2001;18(1):221–7.

17. Peers C, Wyatt CN, Evans AM. Mechanisms for acute oxygen sensing in the carotid body. Respir Physiol Neurobiol. 2010;174(3):292–8.

18. Evans AM, Mustard KJ, Wyatt CN, Peers C, Dipp M, Kumar P, et al. Does AMP-activated protein kinase couple inhibition of mitochondrial oxidative phosphorylation by hypoxia to calcium signaling in O2-sensing cells? J Biol Chem. 2005;280(50):41504–11.

19. Wyatt CN, Mustard KJ, Pearson SA, Dallas ML, Atkinson L, Kumar P, et al. AMP-activated protein kinase mediates carotid body excitation by hypoxia. J Biol Chem. 2007;282(11):8092–8.

20. Dallas ML, Scragg JL, Wyatt CN, Ross F, Hardie DG, Evans AM, et al. Modulation of O(2) sensitive K (+) channels by AMP-activated protein kinase. Adv Exp Med Biol. 2009;648:57–63.

21. Kréneisz O, Benoit JP, Bayliss DA, Mulkey DK. AMP-activated protein kinase inhibits TREK channels. J Physiol. 2009;587(Pt 24):5819-30.

22. Kim D, Kang D, Martin EA, Kim I, Carroll JL. Effects of modulators of AMP-activated protein kinase on TASK-1/3 and intracellular Ca(2+) concentration in rat carotid body glomus cells. Respiratory physiology & neurobiology. 2014;195:19–26.

23. Myers RW, Guan HP, Ehrhart J, Petrov A, Prahalada S, Tozzo E, et al. Systemic pan-AMPK activator MK-8722 improves glucose homeostasis but induces cardiac hypertrophy. Science. 2017;357(6350):507–11.

24. Feng D, Biftu T, Romero FA, Kekec A, Dropinski J, Kassick A, et al. Discovery of MK-8722: A Systemic, Direct Pan-Activator of AMP-Activated Protein Kinase. ACS Med Chem Lett. 2018;9(1):39–44.

25. Sanders MJ, Ali ZS, Hegarty BD, Heath R, Snowden MA, Carling D. Defining the mechanism of activation of AMP-activated protein kinase by the small molecule A-769662, a member of the thienopyridone family. J Biol Chem. 2007;282(45):32539–48.

26. Feng D, Biftu T, Romero FA, Kekec A, Dropinski J, Kassick A, et al. Discovery of MK-8722: a systemic, direct pan-activator of AMP-activated protein kinase. ACS Med Chem Lett. 2018;9(1):39–44.

27. Hardie DG, Schaffer BE, Brunet A. AMPK: An Energy-Sensing Pathway with Multiple Inputs and Outputs. Trends Cell Biol. 2016;26(3):190–201.

28. Ross FA, Rafferty JN, Dallas ML, Ogunbayo O, Ikematsu N, McClafferty H, et al. Selective expression in carotid body type I cells of a single splice variant of the large conductance calcium-and voltage-activated potassium channel confers regulation by AMP-activated protein kinase. The Journal of biological chemistry. 2011;286(14):11929–36.

29. Ikematsu N, Dallas ML, Ross FA, Lewis RW, Rafferty JN, David JA, et al. Phosphorylation of the voltage-gated potassium channel Kv2.1 by AMP-activated protein kinase regulates membrane excitability. Proc Natl Acad Sci U S A. 2011;108(44):18132–7.

30. Moral-Sanz J, Lewis SA, MacMillan S, Ross FA, Thomson A, Viollet B, et al. The LKB1-AMPK-alpha1 signaling pathway triggers hypoxic pulmonary vasoconstriction downstream of mitochondria. Sci Signal. 2018;11(550):eaau0296.

31. Moral-Sanz J, Mahmoud AD, Ross FA, Eldstrom J, Fedida D, Hardie DG, et al. AMP-activated protein kinase inhibits Kv 1.5 channel currents of pulmonary arterial myocytes in response to hypoxia and inhibition of mitochondrial oxidative phosphorylation. J Physiol. 2016;594(17):4901–15.

32. Corton JM, Gillespie JG, Hawley SA, Hardie DG. 5-aminoimidazole-4-carboxamide ribonucleoside. A specific method for activating AMP-activated protein kinase in intact cells? Eur J Biochem. 1995;229(2):558–65.

33. Xiao B, Sanders MJ, Underwood E, Heath R, Mayer FV, Carmena D, et al. Structure of mammalian AMPK and its regulation by ADP. Nature. 2011;472(7342):230–3.

34. Xiao B, Sanders MJ, Carmena D, Bright NJ, Haire LF, Underwood E, et al. Structural basis of AMPK regulation by small molecule activators. Nat Commun. 2013;4:3017.

35. Dite TA, Ling NXY, Scott JW, Hoque A, Galic S, Parker BL, et al. The autophagy initiator ULK1 sensitizes AMPK to allosteric drugs. Nat Commun. 2017;8(1):571.

36. Benziane B, Bjornholm M, Lantier L, Viollet B, Zierath JR, Chibalin AV. AMP-activated protein kinase activator A-769662 is an inhibitor of the Na(+)-K(+)-ATPase. Am J Physiol Cell Physiol. 2009;297(6):C1554–66.

37. Kopietz F, Alshuweishi Y, Bijland S, Alghamdi F, Degerman E, Sakamoto K, et al. A-769662 inhibits adipocyte glucose uptake in an AMPK-independent manner. Biochem J. 2021;478(3):633–46.

